# DoChaP: The Domain Change Presenter

**DOI:** 10.1101/2020.12.16.423045

**Authors:** Shani T. Gal-Oz, Nimrod Haiat, Dana Eliyahu, Guy Shani, Tal Shay

**Affiliations:** Department of Life Sciences, Ben-Gurion University of the Negev, Beer-Sheva, Israel; Department of Software and Information Systems Engineering, Ben-Gurion University of the Negev, Beer-Sheva, Israel

## Abstract

Alternative RNA splicing results in multiple transcripts of the same gene, possibly encoding for different protein isoforms with different protein domains and functionalities. Whereas it is possible to manually determine the effect of a specific alternative splicing event on the domain composition of a particular encoded protein, the process requires the tedious integration of several data sources; it is therefore error prone and its implementation is not feasible for genome-wide characterization of domains affected by differential splicing. To fulfill the need for an automated solution, we developed the Domain Change Presenter (DoChaP), a web server for the visualization of the exon–domain association. DoChaP visualizes all transcripts of a given gene, the domains of the proteins that they encode, and the exons encoding each domain. The visualization enables a comparison between the transcripts and between the protein isoforms they encode for. The organization and visual presentation of the information makes the structural effect of each alternative splicing event on the protein structure easily identified. To enable a study of the conservation of the exon structure, alternative splicing, and the effect of alternative splicing on protein domains, DoChaP also facilitates an inter-species comparison of domain–exon associations. DoChaP thus provides a unique and easy-to-use visualization of the exon–domain association and its conservation between transcripts and orthologous genes and will facilitate the study of the functional effects of alternative splicing in health and disease.

## Introduction

Alternative splicing creates multiple transcripts from a single gene, possibly encoding for different protein isoforms, and therefore increases the functional complexity of the eukaryotic proteome^1–4^. Proteins are composed of structural domains, each of which may have an independent function^5^. Protein domains can be encoded entirely from one exon or from several consecutive exons. In both cases, alternative splicing can cause either the loss of the domain or a change in the length and composition of the RNA sequence that codes for the domain and hence in the functionality of the domain or even of the entire protein. Indeed, it has previously been reported that alternative splicing changes protein functionality in tissue-specific splicing^6^, protein interactions^7^, cancer^8,9^, resistance to biological therapeutics^10^, and the immune response to viral infections^11^.

**Box 1**. Manual determination of the effect of alternative splicing on protein domains in a single gene

1. Identify the transcripts of the gene (from RefSeq and/or Ensembl)
2. Find the protein isoforms that are encoded by the identified transcripts (from RefSeq and/or Ensembl)
3. Predict the domains in each isoform separately (e.g., by Pfam or SMART) or look for the domains of the isoform in databases (e.g., NCBI’s CDD)
4. Manually compare the type, position, and length of the protein domains found between each pair of isoforms.
5. For each domain whose properties have changed, translate the positions of the domain in the two protein isoforms into positions in the transcripts, translate positions in the transcripts into genomic positions, and examine which exon(s) of the transcripts that encode the pair of isoforms are in this position and how this(these) exon(s) has(have) changed between the transcripts.
6. For each domain that is missing in one of the isoforms, translate the position in the protein that carries the domain into the position in the transcript, translate the position in the transcript into the genomic position, and examine which exon is in this position and how this exon has changed between the transcripts.

The use of high throughput RNA sequencing (RNA-seq) data to study genome wide alternative and differential splicing is becoming more and more common^12^. However, determining the functional implications of alternative splicing at the protein level involves a tedious manual process (Box 1) for each case of alternative splicing. Furthermore, experimental biologists who are studying specific cases of different functionalities of alternative protein isoforms currently do not have a tool for exploring the different isoforms of the protein alongside their domains.

Currently available domain prediction tools (e.g., SMART^13^, Pfam^14^, NCBI’s CDD^15^, and TIGRFAMs^16^) predict protein domains either from the amino acid sequence of known proteins or from user supplied query sequences. Such tools provide the domains’ annotation and visualization for the protein of interest, and some even specify the location of the introns^13^, but they can present only a single protein isoform (encoded by a single transcript) at a time, regardless of the other known isoforms and their domain compositions. Several database tools have thus attempted to address the problem of studying the effect of alternative splicing on protein domains, e.g., ExDom^17^, ProSAS^18^, and ASPicDB^19^. However, ExDom^17^ and ProSaS^18^ were built around a decade ago on much smaller source databases, and at present they are down and no longer maintained. ASPicDB^19^ is limited to human genome and does not show the transcript to isoform association or the exon to domain association. Therefore, the gap between the amount of existing information about the effect of alternative splicing on protein domains and the ability of researchers to visualize this effect remains to be breached.

To close this gap and thereby to provide researchers with an intuitive visualization of alternative splicing and information on protein domains, we built the Domain Change Presenter (DoChaP) web server. DoChaP provides a user-friendly, simple and intuitive gene-centric visualization of all the alternative splicing isoforms of a gene and the domains that they encode (Figure 1). DoChaP covers five species and presents the genes in their genomic context, alongside their transcripts and protein domains, thereby highlighting the potential connections between exons and the protein domains that they encode.

**Figure 1.**
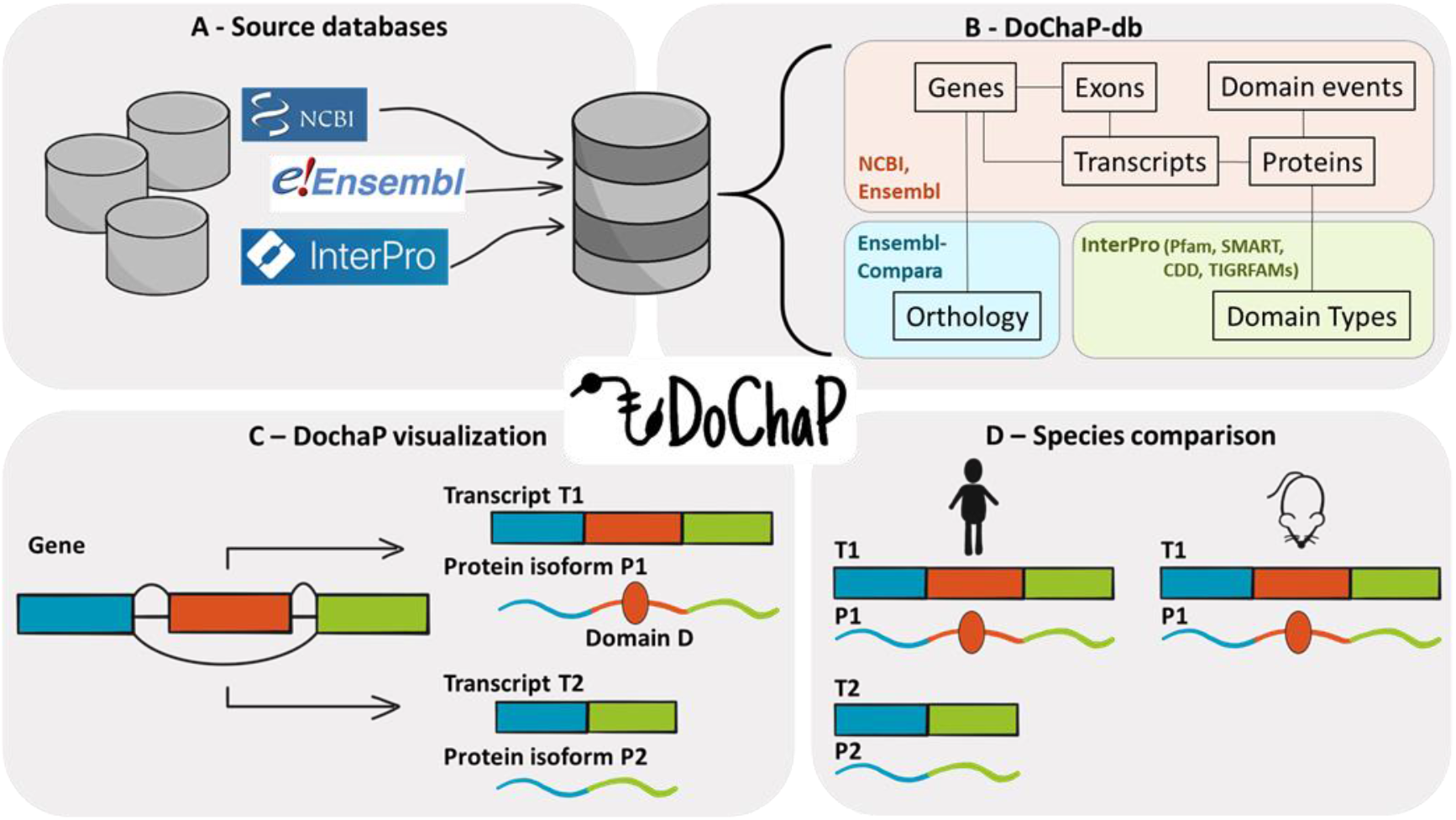
Overview. **(A)** The DoChaP-database (DoChaP-db) integrates information on transcripts and protein domains from several sources into a single SQLite database. **(B)** Genes, transcripts, exons, proteins and domain events data was taken from NCBI and Ensembl. Orthology information was taken from Ensembl Compara. Domain types and the mapping of the domain accessions from different sources (Pfam, SMART, CDD, and TIGRFAMs) was inferred from InterPro. **(C)** A simple visualization of the information collected into the DoChaP-db is presented in the DoChaP webserver for a user selected gene. The visualization presents the genomic context of the transcripts (Gene) and the exon structure of the coding region of the mature mRNA of all the transcripts (T1, T2) and their encoded protein domain composition (P1, P2). Exons and their encoded protein domains are represented in different colors. **(D)** The ‘species comparison’ feature allows the user to select a gene and a species and visualize the orthologous genes so as to enable a comparison of transcripts and their protein domain organization.

## Methods

### Data Sources

DoChaP integrates information for transcripts and protein domains from several sources (Table 1) into a single SQLite database (Figure 1A). RefSeq^20^ and Ensembl’s^21^ transcripts, gene CDSs and exon annotations were extracted from GFF (general feature format) files, downloaded directly from the ftp sites of NCBI’s Genome (https://ftp.ncbi.nlm.nih.gov/genomes/refseq/) and Ensembl (ftp://ftp.ensembl.org/pub/current_gff3/). Protein information, connections between transcripts and protein isoforms, and domain annotations, descriptions and external identifiers were taken from: (1) RefSeq: GenPept flat files (.gpff), downloaded from NCBI’s RefSeq ftp site (ftp://ftp.ncbi.nih.gov/refseq/species-specific/mRNA_Prot/); (2) Ensembl: BioMart data mining tool^22^ (https://m.ensembl.org/biomart/martview/), using xml queries (templates can be found in the GitHub repository; Figure 1B). RefSeq and Ensembl use domain predictions from multiple sources, of which DoChaP presents only NCBI’s Conserved Domains Database (CDD)^15^, Pfam^14^, SMART^13^, TIGRFAMs^16^ and InterPro^23^. For DoChaP, the connection between domain accessions from different sources was inferred from InterPro entries that were downloaded directly from InterPro^23^ (https://www.ebi.ac.uk/interpro/entry/InterPro/#table). The association between RefSeq and Ensembl accession numbers was taken from the Gene2ensembl table downloaded from NCBI’s Gene ftp site (ftp://ftp.ncbi.nih.gov/gene/DATA/; Figure 1B).

**Table 1.**
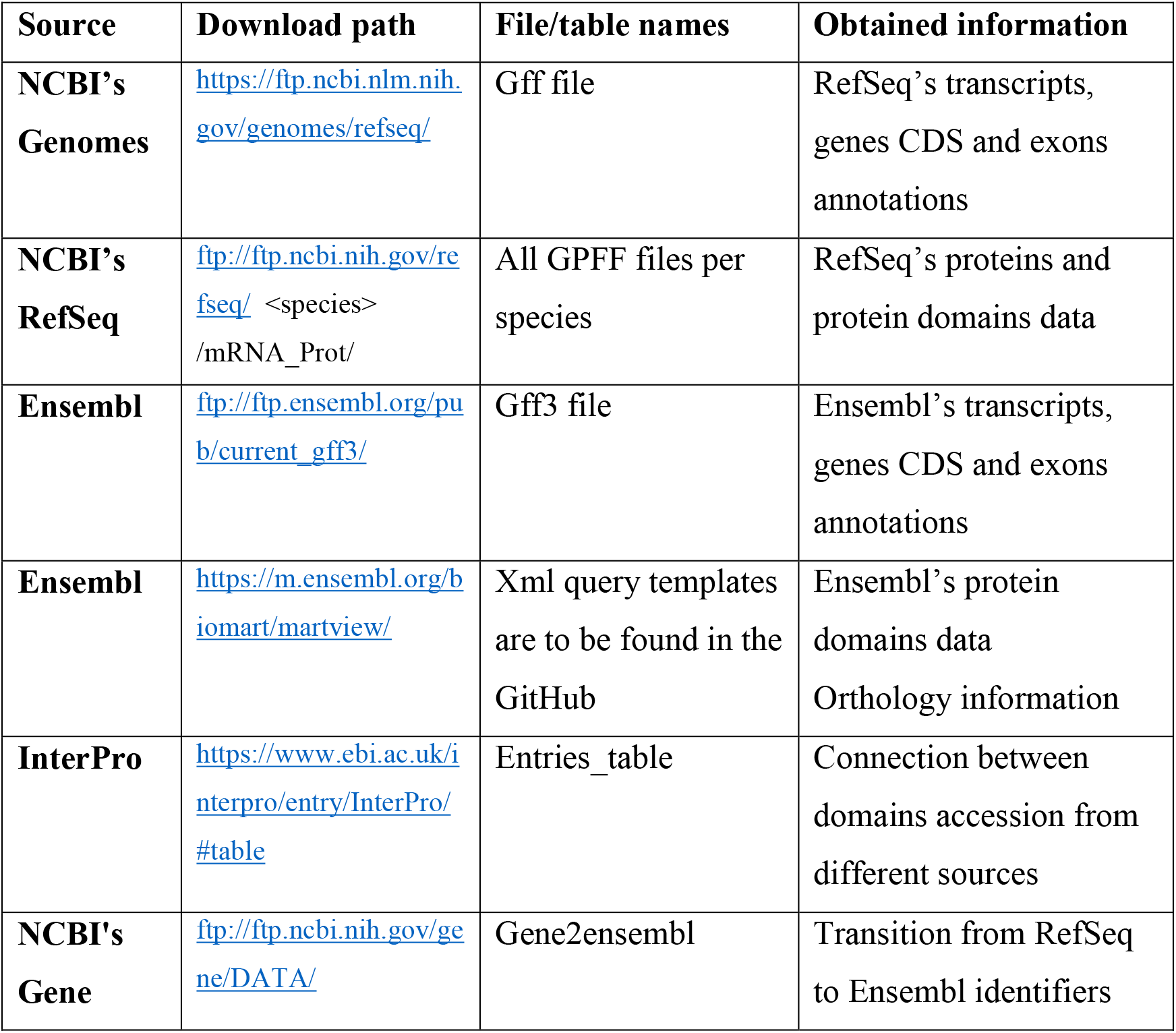
DoChaP sources

Orthology data for each pair of species in the DoChaP database was downloaded from Ensembl Compara via the BioMart data mining tool, using xml queries (xml templates can be found in the GitHub repository; Figure 1B).

### Database content

The DoChaP database currently includes information for five species: human (*Homo sapiens*, hg38), mouse (*Mus musculus*, mm10), rat (*Rattus norvegicus*, rn6), zebrafish (*Danio rerio*, danRer11) and frog (*Xenopus tropicalis*, xenTro9). The database content is detailed in Table 2 (relevant to October 2020), and the database schema is detailed in Supplementary Figure 1.

**Table 2.** DoChaP database content for five species

| Species | Gene IDs | Transcript<br>- isoform<br>pairs | Unique<br>exons | Protein<br>domains | Domain<br>occurrences | In-domain<br>junction |
| --- | --- | --- | --- | --- | --- | --- |
| *Homo sapiens* | 20,388 | 173,783 | 420,565 | 10,855 | 366,414 | 819,397 |
| *Mus musculus* | 22,971 | 118,560 | 351,417 | 10,352 | 267,835 | 581,283 |
| *Rattus norvegicus* | 25,377 | 63,812 | 287,556 | 10,507 | 151,469 | 352,689 |
| *Danio rerio* | 29,538 | 70,105 | 352,955 | 10,165 | 264,619 | 547,192 |
| *Xenopus tropicalis* | 31,912 | 88,166 | 473,954 | 9,277 | 331,223 | 938,047 |
| **Total** | 130,186 | 514,426 | 1,886,447 | 51,156 | 1,381,560 | 3,238,608 |

### Implementation

The DoChaP database was built using Python 3. The Ftplib package (https://docs.python.org/3/library/ftplib.html) was used for downloading from the ftp sites. The Bio.SeqIO module of the Biopython^24^ package was used for parsing the GFF and GPFF; the pandas^25^ package was used for data parsing; and the SQLite3 package (https://docs.python.org/3/library/sqlite3.html) was used for building the database.

The DoChaP web server is implemented with an Express.js on Node.js environment and is distributed via XAMPP. The website is controlled by AngularJS and uses HTML5 canvas for visualization.

## Results

### Features and input

The DoChaP web server currently includes two visualization options for the exon– domain relationship: (1) The ‘single species transcript comparison’ provides genomic, transcript and domain composition visualization of all the known RefSeq and Ensembl transcripts of a gene of interest in a specified species (Figure 1C). (2) The ‘species comparison’ provides the same information for a gene of interest in a selected species and its orthologous gene in one of the other available species (Figure 1D).

The search term for DoChaP can be in any of the following formats: gene, transcript or protein identifiers (gene symbol; NCBI’s Gene ID or RefSeq transcript or protein ID; Ensembl gene, transcript or protein ID). In the single species transcript comparison, the species of interest should be specified. In the species comparison feature, the user should select a gene and species of interest, DoChaP will then search for all the orthologs of the gene in the other species and allow the user to select which species and orthologs to present. Orthologous genes are taken from Ensembl Compara. As some genes with identical gene symbols from different species are not defined as orthologs in Ensembl Compara (e.g., *A2ml1* in human and mouse), comparison of such genes is also enabled in DoChaP.

### Graphical Visualization

DoChaP provides an intuitive, informative and straight-forward way of exploring the annotations of exons and domains. In the results page, all the transcripts of the gene of interest are shown with the following elements:

a. Genomic visualization: The genomic visualization provides the genomic context of the shown transcript, including the loci, strand direction, and introns, exons and coding region. Exons are represented in different colors that are consistent across all the transcripts of the gene and through all the visualizations. Information on the start and end positions of the exon and its ordinal number in the transcript is available by mouse hovering on each exon. Each transcript can be manually hidden from the view by clicking the "hide transcript" button (another click will show it). The user can adjust the genomic range shown so as to focus on a specific set of exons. For genes that are coded on the reverse strand, the genomic visualization will be reversed (horizontally flipped) so that, regardless of the coding strand, the location of first exon is the left-most and that of the last exon is the right-most.
b. Transcript visualization: For each transcript in the genomic visualization, the exon structure of the coding region of the mature mRNA (without the untranslated regions) is provided. Exons are represented in different colors, corresponding to the relevant colors in the genomic visualization and being consistent across transcripts. The length and ordinal number of the exon in the transcript are available by mouse hovering over the exon. Transcript RefSeq and/or Ensembl IDs are shown above each transcript. The nucleotide and amino acid length scales are shown at the top. The user can adjust the genomic range displayed so as to focus on a specific set of exons and domains.
c. Protein composition visualization: Protein domains are shown on a black line below the mRNA exon structure, and are represented with ellipses. The colors and positions of the domains are consistent with the encoding exon or exons. Different colors within the same domain indicate that more than one exon codes for the domain (in-domain junction). The domain name is shown below the domain rounded shape. By mouse hovering over the domain, additional details such as the full name, position on the protein, length in amino acids and link to the external domain source are displayed. For simplicity, overlapping domains are grouped and represented as a single ellipse with a double line. Mouse hovering will show how many domains are represented within this shape and a click on it will expand the visualization to show all the grouped overlapping domains. Protein RefSeq and/or Ensembl IDs are shown above each protein isoform visualization.
d. Color guide exons table: This table is shown on the bottom of the results page. Each color assigned to an exon in that page is shown, and the start and end positions as well as the IDs of all the transcripts in which the exon is included are detailed in the table.

The ‘change display option’ allows the user to choose which types of visualization are shown: genomic only, transcript only, protein composition only or transcript and protein composition together. The default view displays all views. The ‘hide predicted records’ option enables removal of all RefSeq’s predicted transcripts and proteins (prefix XM_ and XP_), and only curated RefSeq (prefix NM_ and NP) and Ensembl (prefix ENS) records will be displayed in the results. In the species comparison tab, all transcripts of the relevant gene from both species are shown in similar format, side by side. Each result page can be exported to a PDF by using the ‘Save as PDF’ button.

### Example and Biological impact

To demonstrate how the visualization of DoChaP can promote the understanding of the functional effect of alternative splicing, we use the tumor suppressor breast cancer gene 1 (*BRCA1*) as a model gene. *BRCA1* is one of the most commonly mutated genes in breast and ovarian cancer^26^. In Figure 2, we present the DoChaP view of human *BRCA1* (4 representative transcripts out of 32). *BRCA1* is encoded on the reverse DNA strand and consists of 24 exons (Figure 2A). Alternative splice variants of *BRCA1*, mainly focused in exons 1, 3 and 10 (E1a/b, E3, E10, respectively; Figure 2B and C), have been previously reported and widely studied in different contexts^27–29^. The frequency of the alternative transcripts of BRCA1 changes between tissues^29^, and also might play a role in the tumorigenesis of breast cancer^30^. For example, exon number 3, is skipped in *BRCA1* transcript NM_007297 (E3, Figure 2A left and zoom in Figure 2B). Since this exon encodes part of the RING finger motif (Figure 2A, right panel), the resulting protein product lacks the RING finger domain (“Znf RING”), and this, in turn, might affect the cell cycle regulatory function of this protein isoform^29^. The largest exon of the *BRCA1* gene is exon E10 (3,426 bp; Figure 2A left and zoom in Figure 2C, light blue). Strangely, this exon is sometimes referred to as exon 11, because of an upstream Alu element that was inserted into BRCA1 clone (historically referred to mistakenly as exon 4)^26^. Two of the presented transcripts (NM_007297 and NM_007300) include the full-length E10 (3,426 bp, Figure 2c, light blue). However, in transcripts NM_007298 and NM_007299, an alternative 5’ splice site is used and only the first 117 bp of this exon (E10117) remain in the mature mRNA (Figure 2c, purple). The alternative 5’ splice site of exon 10 causes the loss of specific protein binding sites [such as RB, p53, MYC, RAD50, TUBG (γ-tubulin) and angiopoietin-1] and the loss of a nuclear localization signal, and therefore affects the functionality of the translated protein^27^. The “BRCA1 serine dom” domain, which is a serine-rich domain associated with the BRCA1 C-terminus, is encoded from the full version of exon 10 (Figure 2a, right panel, light blue) and is seen only in the protein products of the transcripts that include the full-length exon 10 (Figure 2a, right panel). In the case of *BRCA1*, DoChaP provides a clear and intuitive visualization of the complex pieces of information that have been collected over many years in several different studies and of all the known domains that exist (or not) according to the annotation of all transcripts.

**Figure 2.**
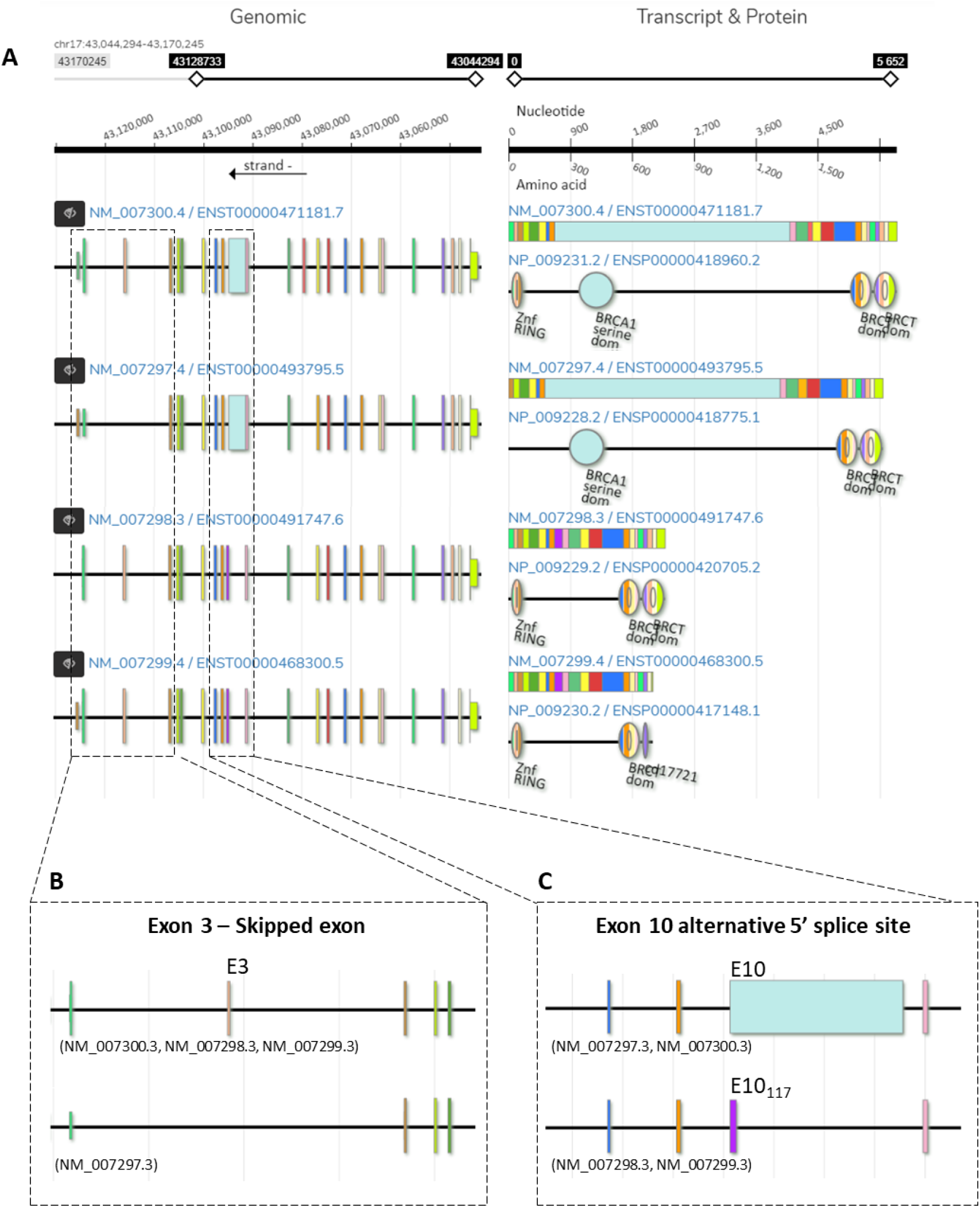
Sample output of DoChaP for the human breast cancer susceptibility gene BRCA1. **(A)** Left, genomic visualization of the transcripts in their genomic context (genomic region, scale and strand are on the top). Right, mRNA and protein domain composition for each transcript. Different colors represent different exons, and are consistent across all the visualizations of the same gene. Domains are shown as circular shapes and are colored according to the exons that encode for them. In BRCA1, the top transcript encodes for four domains. The BRCT domain exists in all four isoforms shown, either once or twice. **(B)** In the second transcript, the third exon (E3) is skipped and therefore the Znf RING protein domains are missing. **(C)** The third and fourth protein isoforms not include BRCA1 serine domain, encoded by exon 10 (E10), as its associated transcripts have a shorter exon 10 (E10117) due to an alternative 5’ splice site event. For the sake of simplicity, only four representative transcripts of BRCA1 are shown.

### Limitations

The public databases used as sources for DoChaP include redundancies and inconsistencies. To overcome this problem, we currently use the RefSeq – Ensembl conversion table gene2ensembl (from NCBI’s ftp, see methods). In cases where matched RefSeq and Ensembl protein records have different lengths (of more than one amino acid; for example, the protein isoform NP_001269863.2 and ENSP00000408592.1 of the gene Prdm15 have different lengths of 1,161 and 1,198 amino acids, respectively), the records were separated into two transcripts in the visualization. In all other cases, matched records were merged into one transcript to avoid redundancies.

Different data sources may indicate slightly different start and end positions of the domains in the protein, probably due to differences in domain prediction methods [for example, in the protein isoform NP_000236.2 of the human gene Met, pfam has located the Sema domain between amino acids 57-490 (pfam01403) and SMART has located the same domain in 52-496 (smart00630)]. In such cases, we present the coordinates of all the overlapping domains and show the source from which the coordinates of each domain occurrence were taken. Overlapping domains are shown as one ellipse with inner circle line that can be expanded upon click.

Some very long transcripts and transcripts with many exons make the genomic visualization of an entire gene less efficient in terms of colors and relative exon sizes. In such cases, the user is advised to zoom in and explore the gene in several consecutive windows.

## Conclusions and Future Extensions

Improved sequencing technologies are leading to increased interest in the connection between exons and domains and the evolution of the regulatory mechanisms of alternative and differential splicing. Previously, to explore exon–domain associations, researchers had to manually search, integrate and interpret textual data from several sources in a tedious and error-prone process. With the development of DoChaP (made possible by the increasing quantities of freely available data in the public domain), the research community has access to a fast, intuitive and easy-to-use visualization tool for exploring the exon–domain relationship.

DoChaP is an ongoing project, and we plan to add new features and visualization capabilities and increase the database in updates. Specifically, we intend to add additional species and other types of functional features of proteins, such as signal peptides. Moreover, we will integrate a method for comparison of transcripts that will allow DoChaP to order the transcripts based on their genomic similarity and domain organization and to present similarity scores for transcripts of the same gene and between orthologs.

Given the increased interest in the effect of alternative splicing on functionality in health and disease, we believe that DoChaP will allow researchers from the biological, medical and computational fields to interpret the structural effect of their transcriptional profiling changes toward new functional findings.

## Data Availability

The DoChaP web server is freely available at https://DoChaP.bgu.ac.il. The latest DoChaP database may be downloaded under “Downloads” tab in the DoChaP website. The source code is available on GitHub (https://github.com/Tal-Shay-Group/DoChaP).

## Acknowledgements

The authors would like to thank Yehuda Barouch for his continuous valuable support during DoChaP server development, and Eran Lachs and Yossi Gross from the Technologies, Innovation & Digital Division at Ben-Gurion University for the technical support during server setup.

