## Supplementary Figure 1 for "DoChaP: The Domain Change Presenter"

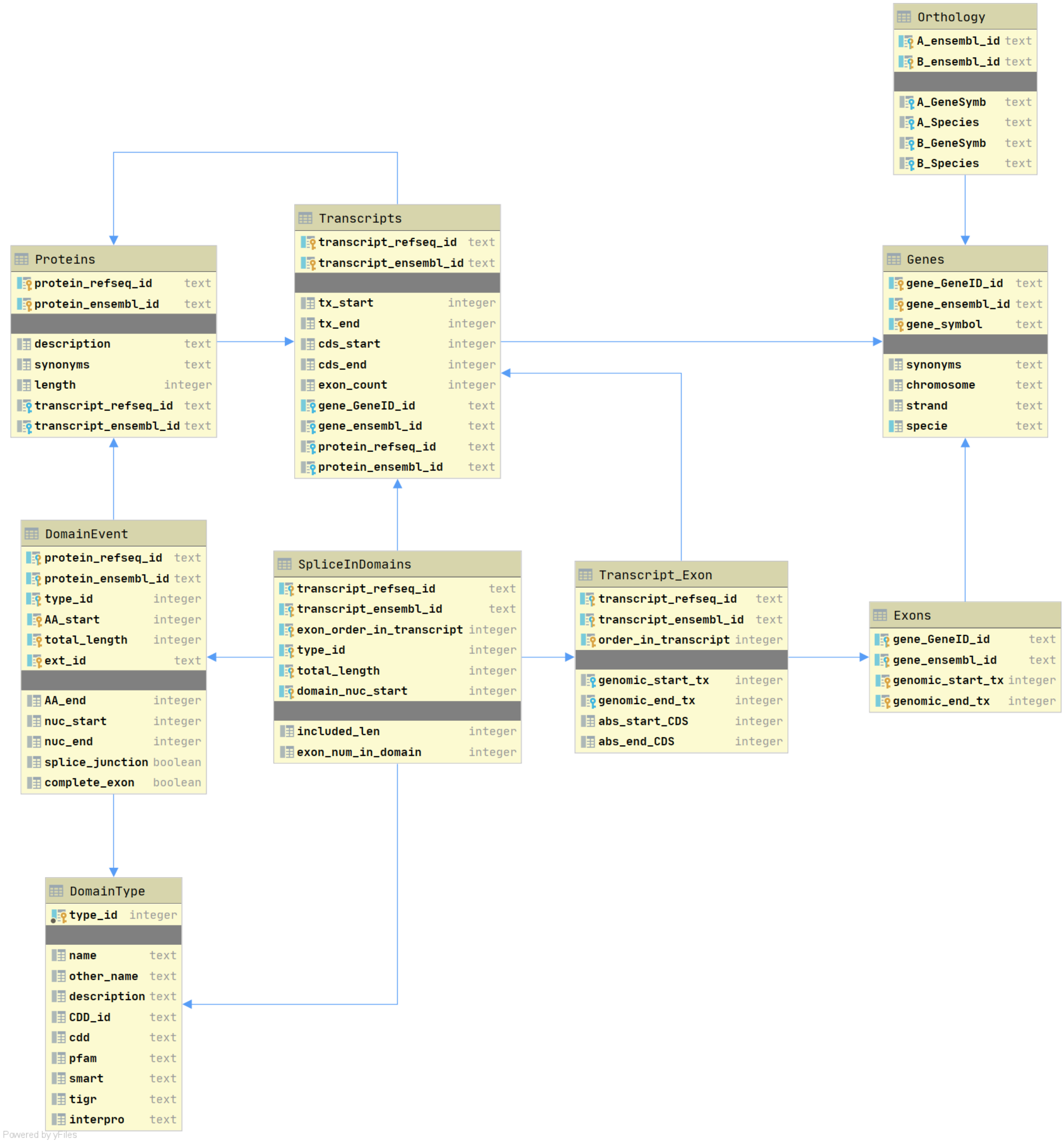


**Supplementary Figure 1. DoChaP database schema.** DoChaP database (DoChaP-db) is a SQLite database, in which: (1) For each gene, all the exons, protein coding transcripts and their corresponding protein isoforms are listed (tables: Genes, Exons, Transcripts, and Proteins). (2) For each transcript, all the exons are detailed (table: Transcript_Exon). (3) For each protein isoform, all domain occurrences are listed (in table DomainEvent), where each domain occurrence is a specific case of a domain type (listed in table DomainType). (4) Splice junctions that occur within a domain coding sequence are termed in-domain junctions (listed in the table SpliceInDomain). (5) All pairs of orthologous genes between species in the database are listed in the table Orthology.
